# Trade-offs of managing Arctic predator harvest instability in fluctuating oceans

**DOI:** 10.1101/2020.06.17.154971

**Authors:** Daisuke Goto, Anatoly A. Filin, Daniel Howell, Bjarte Bogstad, Yury Kovalev, Harald Gjøsæter

## Abstract

Sustainable human exploitation of marine living resources stems from a delicate balance between short-term yield stability and long-term population persistence to achieve socioeconomic and conservation goals. However, imperfect knowledge of how oscillations in ecosystem processes regulate fluctuations in exploited populations can obscure the risk of missing management targets. We illustrate how the harvest policy to suppress short-term yield fluctuation inadvertently disrupts population cycles and yield stability of exploited, long-lived predators under stochastically fluctuating environmental forces (food availability and regional climate) using Northeast Arctic (NEA) cod (*Gadus morhua*, an apex predatory fish) as a case study. We use a stochastic, empirically parameterized multispecies model to simulate NEA cod population dynamics through life-history processes; Barents Sea capelin (*Mallotus villosus*, a pelagic forage fish) modulates cod productivity through density-dependent cannibalism–predation dynamics, whereas sea temperature regulates cod consumption, growth, and recruitment. We first test how capelin and sea temperature fluctuations regulate patterns in cod yield fluctuation and then quantitatively assess how fishing pattern designed to limit yield between-year variance (within 50–5%) perturbs cod population–catch dynamics. Simulations suggest that capelin and temperature interactively contribute to shifting cyclic patterns in cod yield fluctuation primarily through cod cannibalism–predation dynamics. Wavelet analyses further show that muffling yield variance (30 % or less) reshapes the cyclicity (shorter period and greater amplitude) of cod population size and demography, thereby becoming progressively unsynchronized with fishing pressure. Our work reveals unintended consequences of managing transient dynamics of fished populations: the interworking of population cycle destabilized by inadvertently intensifying fishing pressure, amplifying yield fluctuation and, in turn, elevating overharvest risk when not accounting for compounded effects of stochasticity in ecologically connected processes. These policy implications underscore the need for an ecosystem approach to designing ecologically sound management measures to safely harvest shared living resources while achieving socioeconomic security in increasingly more dynamic oceans in the Arctic and elsewhere.

## INTRODUCTION

Fluctuation in wild animal populations has posed myriad socioeconomic and conservation challenges in sustainably managing human exploitation of living natural resources for centuries (Hjort 1914, Fryxell et al. 2010, Carpenter et al. 2015). In commercial exploitation of oceanic species such as long-lived, apex predators, stable population dynamics provide not only predictability to harvesters and seafood processors but also food, nutrition, and employment security in fishing nations (Garcia and Rosenberg 2010, Hicks et al. 2019). In modern industrial fisheries, harvest strategies are designed to maintain population abundance of exploited species above a threshold that prevents overharvest and sustains its harvest while suppressing yield fluctuation to achieve socioeconomic goals (Caddy and Seijo 2005, Nielsen et al. 2018). However, oscillations in environmental forces may amplify or dampen fluctuation in fished populations (Shelton and Mangel 2011, Rouyer et al. 2012). Further, the demography of these populations has often been modified by long-term perturbations such as fishing-induced selective reduction in adult survival (Hidalgo et al. 2011). Harvest strategies that fail to account for the dynamic interplay of environmental and demographic fluctuations may, thus, lead to exploitation intensity that is unsynchronized with natural population cycles, potentially risking sustainability of populations and fishing operation (Hall and Mainprize 2004, Carpenter et al. 2015) and, in turn, incurring ecological and socioeconomic damages (Caddy and Seijo 2005, Nielsen et al. 2018).

Population fluctuation in oceanic predators can arise from stochasticity in demographic states that reflect life-history responses (such as reproductive and migration timing) to large-scale climate variability–an overarching force that governs dynamic habitat conditions (temperature, pH, dissolved oxygen, etc.) over varying spatial and temporal scales (Bjørnstad and Grenfell 2001, Ottersen et al. 2001, Cury et al. 2008). These dynamics often propagate through species interactions to modulate productivity of fished populations (Ottersen et al. 2001, Stige et al. 2010). Environmentally driven plankton production, for example, regulates forage fish production that support exploited apex predators (Ware and Thomson 2005). Large between-year fluctuation can emerge particularly in early life stages of marine populations (Leggett and Deblois 1994); seasonal to decadal variation in ocean conditions propagates through predator–prey interactions, amplifying or attenuating variance in year-class strength of exploited predators (Edwards and Richardson 2004). Stochasticity filtered through such interactions can, thus, emerge as complex, nonlinear patterns in population cycles (Bjørnstad and Grenfell 2001) and pose a challenge in designing management measures to suppress yield variance in exploited predators (Essington et al. 2018).

Trophic interactions between cod (*Gadus morhua*) and capelin (*Mallotus villosus*) in the North Atlantic present such dynamics that modulate short-term fluctuation in cod fishery yield (Link et al. 2009). Productivity of cod–a long-lived, demersal (bottom-water) predatory fish–often fluctuates from year to year with variability in both density-dependent and - independent ecosystem (bottom-up) processes including interactions with other species during early life stages (Lilly et al. 2008, Minto and Worm 2012). In particular, food shortage can enhance cannibalism intensity (Yaragina et al. 2009). Capelin–a short-lived (≤ 5 years), semelparous, widely distributed pelagic planktivore–comprises a large part of adult cod diet (Holt et al. 2019) and may influence the intensity of cod cannibalism on the young (≤ 3-year-olds) and, in turn, recruitment pulses. Moreover, owing to migratory behaviors driven by ocean conditions and plankton distribution, time-varying capelin aggregation may reflect changes in ecosystem processes including systemwide shifts in ice phenology, thermal regimes, and plankton productivity (Carscadden et al. 2001, Rose 2005). Cod–capelin spatial and temporal overlap in cod spawning and nursery grounds may, thus, modulate cod population and yield fluctuations (Ciannelli and Bailey 2005, Johannesen et al. 2012). It remains unclear, however, how fluctuations in bottom-up processes such as forage fish production shape the population behavior and yield stability of intensely exploited predators such as cod, which are concurrently regulated by annual fishing pattern designed to achieve management goals.

Here we illustrate how the harvest policy of suppressing yield fluctuation can reshape the population cycle of exploited predators in stochastically fluctuating environments using a well-studied species, Northeast Arctic (NEA) cod, as a real-world case study. NEA cod, distributed over the Barents Sea and its surrounding areas (Fig. 1a), is the world’s largest supporting two major fishing nations, Norway and Russia (Kjesbu et al. 2014). The fishery dates back to the period when the northern coasts of these nations were likely populated after the end of the last ice age. An international industrial trawl fishery in the Barents Sea developed in the early 1900s and total catches by all gears peaked at more than 1.2 million t in catch in some years in the period 1955–1975 (ICES 2019). Following the establishment of 200-mile economic zones in the late 1970s, the fishery by Norway and Russia dominated (ICES 2019) and subsequently depleted the population prior to the establishment of a harvest strategy by the Norwegian-Russian Fishery Commission in the early 2000s (ICES 2019). This harvest strategy (which was in force until 2016) sets target exploitation rate with precautionary measures while achieving short-term yield stability such as limiting between-year catch fluctuation (to less than 10%). These harvest rules, along with favorable habitat quality (Ottersen and Stenseth 2001, Stige et al. 2010), are suspected to allow the population to balloon with the quota reaching over one million t in 2014, the highest in more than three decades (Kjesbu et al. 2014).

**Figure 1.**
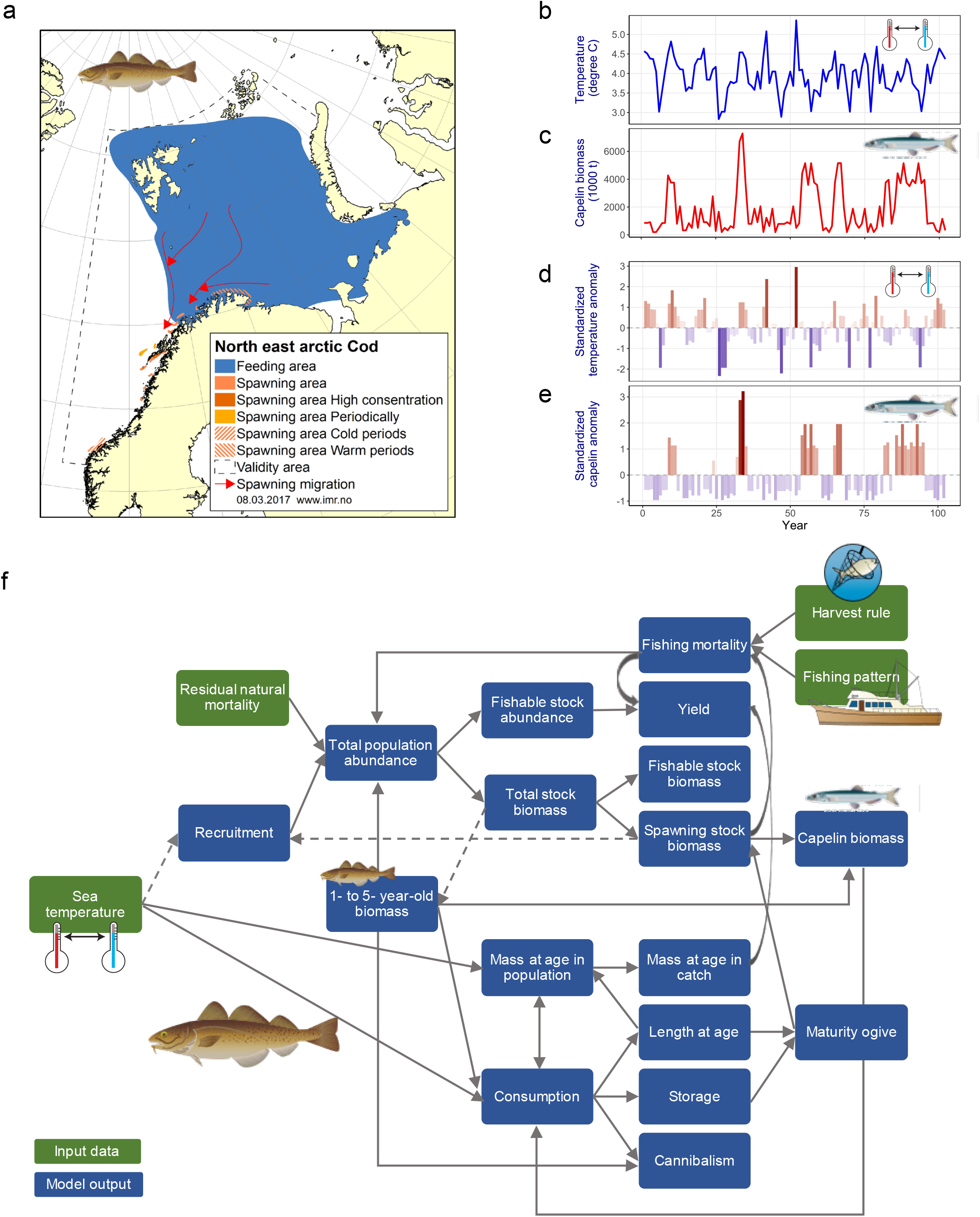
Study system and food-web model. (a) Northeast Arctic (NEA) cod distribution in the Barents Sea. (b) Example of modeled sea temperature (°C) time series based on observed annual mean temperatures in the Kola section. (c) Example of modeled capelin biomass (1000 t) time series. (d) Standardized sea temperature anomalies computed from modeled sea temperatures. (e) Standardized capelin anomalies computed from modeled capelin biomass. (f) Schematic diagram of the NEA cod–capelin stochastic, multispecies model (STOCOBAR) structure. In d and e, standardized anomalies were computed by subtracting modeled annual temperatures (b) or capelin biomass (c) from the mean and dividing by the standard deviation. Artwork: Courtesy of the Integration and Application Network, University of Maryland Center for Environmental Science (ian.umces.edu/symbols/).

Using a stochastic, process-based multispecies model previously developed for the Barents Sea cod–capelin complex (Filin 2009, Howell et al. 2013), we test 1) how climatic oscillation propagates through an interplay of cod cannibalism–capelin productivity (Hjermann et al. 2007) and 2) how fishing pattern to suppress cod yield fluctuation shapes the cyclicity of population size and demographic fluctuations and, in turn, the potential to achieve management targets in stochastically fluctuating environments. By using modeled oscillatory fluctuations in sea temperature in the Kola section (Fig 1b, d)–an indicator for reginal climate (Ingvaldsen et al. 2003) and capelin productivity (Fig. 1c,e)–proxy for the ecosystem carrying capacity (Gjøsæter 1998), in effect, our work aims to quantitatively evaluate implications of adapting an ecosystem approach to provisioning of annual catch targets to ensure sustainable exploitation of marine resources while conserving ecological integrity under fluctuating ocean conditions.

## METHODS

### Study system

The Barents Sea cod–capelin complex is among the few co-managed stocks in the North Atlantic (Gjøsæter et al. 2012). Historically, capelin has been the major forage species supporting exploited apex predators in the Barents Sea food web (Gjøsæter 1998); its stock biomass has been measured by acoustic methods to reach up to seven million t in the 1970s and 1990s (Gjøsæter 1998). Capelin–a small, pelagic, schooling fish, rarely reaching lengths above 20 cm and ages above five-year-olds–feeds in the northern Barents Sea on meso- and macrozooplankton but migrates to the northern coasts of Norway and Russia to spawn in spring. The forage fish is semelparous and thus magnifies its role of bringing energy from the northern parts of the Barents Sea to the southern coastal areas. Although capelin supplies a food source for many piscivores in the Barents Sea, Northeast Arctic (NEA) cod are by far their primary predator, consuming up to more than three million t per year (Dolgov 2002). Capelin fluctuation may, thus, regulate the intensity of cod cannibalism and year-class strength, likely contributing to depensatory effects of depleted food supply (Hjermann et al. 2007). For example, amplified year-to-year fluctuations in capelin biomass in the 1980s and 1990s were largely responsible for the NEA cod biomass rebuilding; cod cannibalism peaked following capelin collapses (Hjermann et al. 2007). The collapses, particularly the first one, propagated through the rest of the Barents Sea food web to a variable degree, depending on availability of other food (Hjermann et al. 2007, Gjøsæter et al. 2009).

### Food-web modeling

To simulate the Barents Sea cod-capelin dynamics, we used a stochastic, food-web model that mechanistically accounts for ecosystem processes (a model of intermediate complexity for ecosystem, Plagányi et al. 2014)–STOCOBAR (Stock of Cod in the Barents Sea). The model has been developed previously at Russian Federal Research Institute of Fisheries and Oceanography (Filin 2009); the version used in this study has been described fully in Howell et al. (2013) (Fig. 1f). For brevity, here we outline the model structure; STOCOBAR is designed to capture age-structured population dynamics of NEA cod in Barents Sea and adjacent areas through yearly life history processes: growth, feeding, maturation, recruitment, natural mortality (including cannibalism on juveniles), and fishing mortality (by single aggregate fleet fishing operation) (Howell et al. 2013). Individual fish growth is mechanistically simulated through energetics by converting consumed food to body mass, which, in turn, regulates maturation (Howell et al. 2013). The model projects cod population dynamics and evaluates alternative management strategies (harvest measures and recovery plans) by accounting for variability in ecosystem processes such as lower trophic group (one or more prey) productivity (Howell et al. 2013). In this study, we incorporated stochasticity explicitly in cod recruitment, capelin productivity, and sea temperature. We parameterized the model with 30-year (1981–2010) annual bottom trawl and acoustic survey data of age-specific life-history traits including maturity rates (proportions of adults) and body masses (kg), and fit it to time series of cod age-specific abundance indices (catch per unit effort) and commercial fishery catch (t), and capelin biomass (t), which were all taken from annual assessments by the International Council for the Exploration of the Sea (ICES) Arctic Fisheries Working Group; the survey and sample processing methods used are extensively documented in ICES (2010).

Cod recruitment (survival of 3-yr-olds) in the model is stochastically simulated and depends directly on 1-year-old cod abundance and cannibalism and indirectly on capelin biomass and sea temperature. We simulated 1-yr-old cod abundance using a density- and temperature-dependent function relating it to spawner abundance (spawning stock biomass, SSB, t) with the Ricker model (Ricker 1958) and mortality by cannibalism (before recruitment). SSB is computed by multiplying age-specific numbers by age-specific masses and maturity rates. The strength of cod cannibalism fluctuates over time and was modeled with a function of capelin biomass and cod population structure (size and age composition) (Fig. 1f, Howell et al. 2013). Although sea temperature may affect cannibalism intensity indirectly through consumption, we assumed that this effect is negligible within the historical temperature range used in this study. Further, we did not include indirect temperature effects on cannibalism through spatial distribution (Howell and Filin 2014). Based on model fitting, we capped mortality by cannibalism at 90%, 60%, and 40% of total mortality of 1-, 2-, and 3-yr-olds (respectively). We then added random noise to recruitment using normally distributed residuals generated from the 1980–2007 hindcast simulations (Howell et al. 2013).

### Modeling capelin and sea temperature fluctuations

We used capelin biomass as proxy for cod carrying capacity, representing “good” and “poor” years of habitat conditions that regulate cod growth and cannibalism. The model uses capelin as a primary food source for cod with all other prey (young Northeast Arctic haddock *Melanogrammus aeglefinus*, young herring *Clupea harengus*, polar cod *Boreogadus saida*, shrimp *Pandalus borealis*, and krill Euphausiidae) being aggregated as “other food’. We statistically modeled between-year variability in capelin production (as an aggregate of 1-to 5-yr-olds) to capture observed patterns of declining biomass 1) when cod SSB exceeding 400,000 t or 2) when capelin biomass falling below three million t in the previous year (no feedback for cod predation on capelin), while capelin biomass is capped at six million t when cod SSB exceeds 800,000 t. Modeled capelin biomass stochastically varies between years and simulations (Fig. 1c), generating low-frequency oscillations (Fig. 1e)

Sea temperature regulates cod population dynamics through recruitment, growth, and consumption to capture variable metabolic rate in the model (Fig. 1f). In this study, we primarily focused on short-term sea temperature fluctuations (rather than multidecadal shifts), a time scale relevant for provisioning of tactical advice in fisheries management. The model uses a stochastically simulated thermal regime that reflects the historical fluctuation of annual mean temperatures (AMT) in the Kola Section in the Barents Sea, which also roughly represents climate variability in the region (Ingvaldsen et al. 2003). To create a temperature anomaly scenario, we first split the observed AMT data (1950-2017) into three groups; cold (AMT < 3.6°C), moderate (3.6°C ≤ AMT ≤ 4.2°C), and warm (AMT > 4.2°C) years. Then, AMT data were randomly selected from these three groups sequentially. The modeled thermal periods (cold, moderate, and warm) last from one to five years; the length of each period was set randomly (effectively generating roughly decadal oscillations, Fig. 1b,d).

### Cod harvest management model and scenarios

We simulated dynamic cod mortality by fishing with a harvest management model based on the harvest rule set for NEA cod (prior to changes in 2016, ICES 2016) and evaluated cod population dynamics and between-year yield variability under alternative harvest scenarios. Under the harvest rule, target exploitation rate (*F_target_*) is set to 0.40 to project annual fishery catch (or total allowable catch, TAC) when spawner abundance (SSB) remains above 460,000 t. These reference values (*F_pa_* and *B_pa_*, respectively) are designed to take precautionary measures to prevent overexploitation by accounting for uncertainty in population and harvest dynamics (Rindorf et al. 2016). When SSB falls below *B_pa_*, however, *F_target_* is adjusted to *F_pa_* scaled to the proportion of SSB relative to *B_pa_* as:

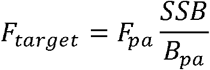

In simulations, *F_target_* was applied by computing age-specific fishing mortality rate as:

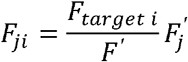

where *F_ji_* is instantaneous rate of fishing mortality of *j*-yr-olds in year *i*; *F_target i_* is target exploitation rate in year *i*, computed in the model according to the harvest rule; *F*’ is mean fishing mortality rate of adults (a fixed age range, 5-to 10-yr-olds, is used for this computation) in year 1; and *F’_j_* is fishing mortality rate of *j*-yr-olds in year 1.

The harvest policy to suppress yield fluctuation (hereafter stability constraint) is designed to provide stable and predictable catch forecasts (and revenue) for harvesters and seafood processors. We computed between-year variation in catch forecasts (or interannual catch variability, *IAV*) as

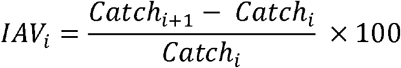

where *IAV_i_* is a relative change in catch forecast (%) from year *i* to *i*+1; *Catch*_*i*+1_ and *Catch_i_* are catch forecasts in year *i*+1 and *i* (respectively). However, the precautionary principle also applies to this policy; the constraint is applied only when SSB equals to or exceeds *B_pa_*. In this study, if this rule results in *IAV* exceeding the set proportion (±%), the model algorithm searches for *F_target_* until it finds the value that does not exceed the boundary set by the constraint. The constraint was not applied in the first year after the recovery period (when SSB ≤ *B_pa_*); *F_target_* was set to *F_pa_* regardless of catch forecasts in the previous year.

To first evaluate how ecological uncertainty involving capelin and climate variability contributes to between-year variance in cod catch forecasts under stability constraint, we tested performance of the following four model configurations that represent ecological assumptions on cod predation–cannibalism dynamics under a 10% constraint with variable sea temperature: M1) constant capelin biomass (one million t) without cannibalism (temperature fluctuation only), M2) varying capelin biomass without cannibalism, M3) constant capelin biomass with cannibalism, and M4) varying capelin biomass with cannibalism.

To evaluate how suppressing short-term variation in catch forecasts drives cod population cycles and yield stability under variable capelin productivity and sea temperature, we analyzed the following nine harvest scenarios of limiting between-year variation in catch forecasts; ±0%, ±5%, ±10%, ±15%, ±20%, ±25%, ±30%, ±40%, and ±50%.

We projected 100-year population and harvest dynamics to evaluate ecological assumptions and harvest scenarios (25 x (4+9) simulations). We analyzed each scenario based on 25 realizations because preliminary simulations showed that mean catches vary less than ±1% with more than 25 simulations. We disregarded the first 20 yrs of model projections as a ‘burn-in’ to minimize effects of the initial conditions and evaluated the outputs from years 21–100. Because this study primarily focuses on implications of suppressing yield fluctuations, we did not incorporate other sources of uncertainty such as population estimation error and implementation error (annual yields were assumed equal to catch forecasts). We set fishing selectivity (age-dependent gear selectivity) as time-invariant in the first year. For clarity, we excluded the following harvest rules set for NEA cod; setting catch forecasts for three years by applying target exploitation rate to current SSB (“three-year forecast”), fixing the lower range of *F_target_*, and raising *F_target_* when SSB becomes exceptionally high (ICES 2019). In analyzing the harvest scenarios, we computed the following six cod demographic state variables and management targets from model projections: harvestable (3-yr-olds and older) population biomass (stock size, t), SSB (t), recruitment, catch forecasts (t), mean fishing mortality, and risk (%). Risk is defined as the probability of SSB falling below the biological limit threshold (*B_lim_*). When SSB < *B_lim_*, the reproductive capacity of a population is expected to decline (Rindorf et al. 2016); harvest rules with risk > 5% are considered unacceptable (non-precautionary) for sustainable exploitation. For NEA cod, *B_lim_* is set to 220,000 t (ICES 2016).

### Spectral analyses

To explore potential mechanisms underlying oscillatory behaviors of projected cod population and yield fluctuations, we performed wavelet spectral analysis on model outputs from select scenarios of environmental forces (sea temperature and capelin) and harvest strategies (±0%, ±10%, ±20%, and ±30% stability constraints). Wavelet spectral analysis is suited for analyzing nonlinear and non-stationary time series (Cazelles et al. 2014) such as population dynamics of exploited species. We computed continuous Morlet wavelet power (the square of amplitudes) spectra to transform signals in the projected time series (years 21–100) as a function of frequency (periodicity) and time from 1000 bootstrap simulations (randomized surrogate time series) (Roesch and Schmidbauer 2018). We derived times series of the following five cod demographic state variables and management targets; stock size, SSB, recruitment, catch forecasts, and mean fishing mortality. Further, to assess (a)synchronicity between cod population and harvest dynamics, we performed cross-wavelet analysis to test for coherency (covariation in frequency) (Cazelles et al. 2014) between stock size and mean fishing mortality time series. All wavelet analyses were performed in R (version 3.6.2, R Development Core Team 2019) with the WaveletComp R package v.1.1 (Roesch and Schmidbauer 2018).

## RESULTS

### Ecological uncertainty in cod harvest dynamics

Under the baseline scenario, cod population dynamics reached dynamic equilibrium by year 20 with, on average, ~2.3 million t harvestable (3-yr-olds and older) biomass (stock size),~1.1 million t spawners (SSB), 694 million recruits (3-yr-olds), and 680,000 t/yr catch (Table 1 and Appendix 1: Fig. S1), which all aligned closely with the historical population status and harvest of NEA cod (ICES 2017). Overall, between-year variability in cod body masses, maturity rates, and consumption rates declined with age (from ~39% to ~7%, Appendix 1: Fig. S1 and S2). Cod diet, on average, comprised of 10% juvenile cod (more than 66% as 1-yr-olds), 21% capelin, and 69% others (Appendix 1: Fig. S2).

**Table 1.**
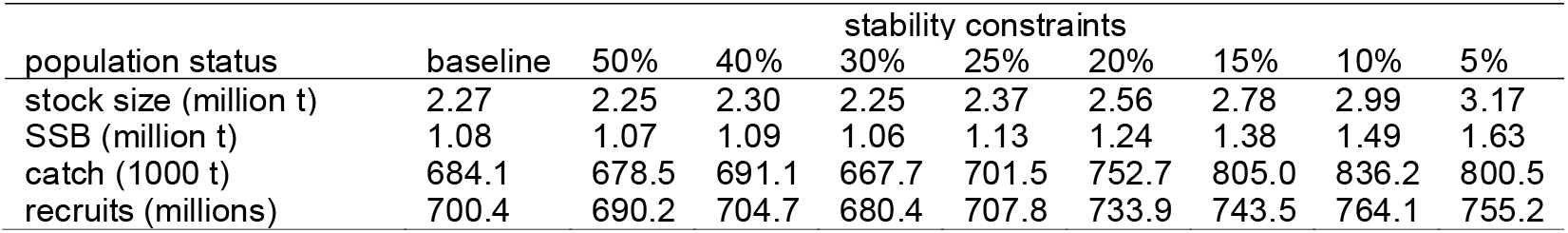
Summary of Northeast Arctic (NEA) cod population status and management target metrics computed from stochastic, multispecies model (STOCOBAR) simulations under scenarios of varying levels of limiting yield fluctuation (stability constraints). Stock size indicates total harvestable (3-year-olds and older) biomass; SSB indicates spawning stock biomass; recruits indicates abundance of 3-year-old cod.

Incorporating additional ecological reality (time-varying capelin productivity and cod cannibalism) in the model showed nonstationary patterns (varying amplitude and periodicity) of cod catch forecasts under a 10% constraint with varying sea temperature (Fig. 2). With constant capelin productivity and no cod cannibalism (M1), wavelet analyses showed a ~8-yr cycle with modest amplitudes in catch forecasts (reflecting one of dominant sea temperature signals, Fig. 2a and Appendix 1: Fig. 3a) with between-year variation of less than 2%/yr (Fig. 2e,f). With varying capelin productivity (M2), catch forecasts showed two (8 to 10- and 32-yr) weaker cycles (roughly reflecting dominant capelin signals, Appendix 1: Fig. 3b), and between-year variation rose only by ~4% (Fig. 2b and 2e,f). By contrast, when cod cannibalism was introduced with constant capelin productivity (M3), catch forecasts showed a strong, consistent 8-yr cycle and substantially higher (–56% to 70%) between-year variation (Fig. 2c and 2e,f). When combined with varying capelin productivity (M4), however, catch forecasts showed an 8-yr cycle with slightly weaker amplitudes and highly erratic between-year variation (ranging from –90% to 312%, Fig. 2d and 2e,f).

**Figure 2.**
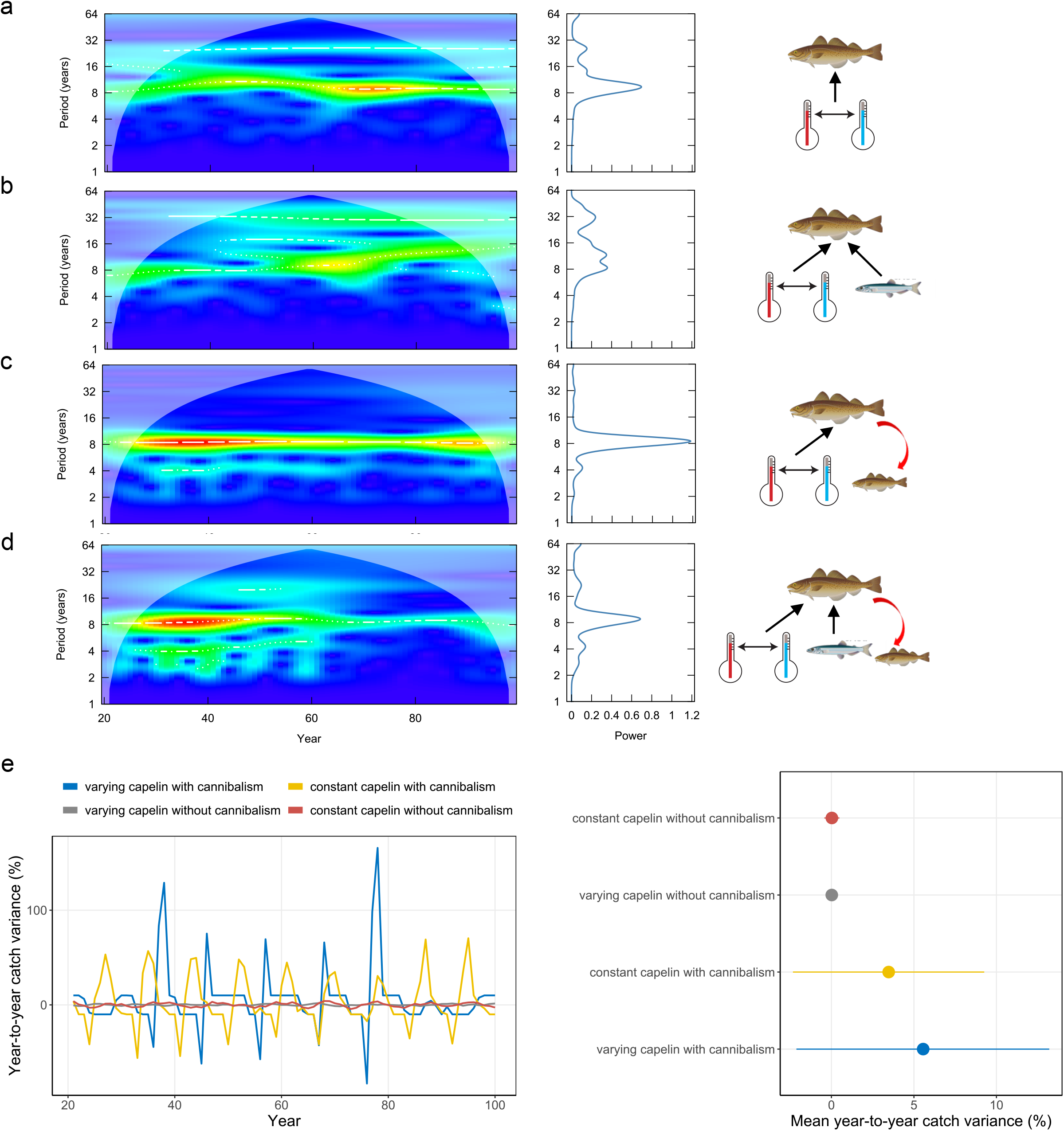
Ecological uncertainty in Northeast Arctic (NEA) cod catch fluctuation projected with a stochastic, multispecies model (STOCOBAR) under scenarios of a ±10% stability constraint and variable sea temperature with varying model configuration of cod cannibalism–capelin productivity. (a–d) Wavelet power spectra [local (times series, left) and global (time-averaged, right)] of NEA cod catch forecasts with (a) constant capelin biomass (1000 t) without cannibalism, (b) varying capelin biomass without cannibalism, (c) constant capelin biomass with cannibalism, and (d) varying capelin biomass with cannibalism. (e) Time series (years 21–100, left) and mean year-to-year variance (%) of annual NEA cod catch (right). In wavelet spectra, red areas indicate higher power (intensity of periodicities), blue areas indicate lower power, and white areas indicate regions influenced by edge effects (outside the “cone of influence”) and inferences cannot be made (Cazelles et al. 2014). In a– d, y-axis is in the logarithms to the base 2. In e (right), error bars indicate 95% confidence intervals (among-year variation over years 21–100). Artwork: Courtesy of the Integration and Application Network, University of Maryland Center for Environmental Science (ian.umces.edu/symbols/).

**Figure 3.**
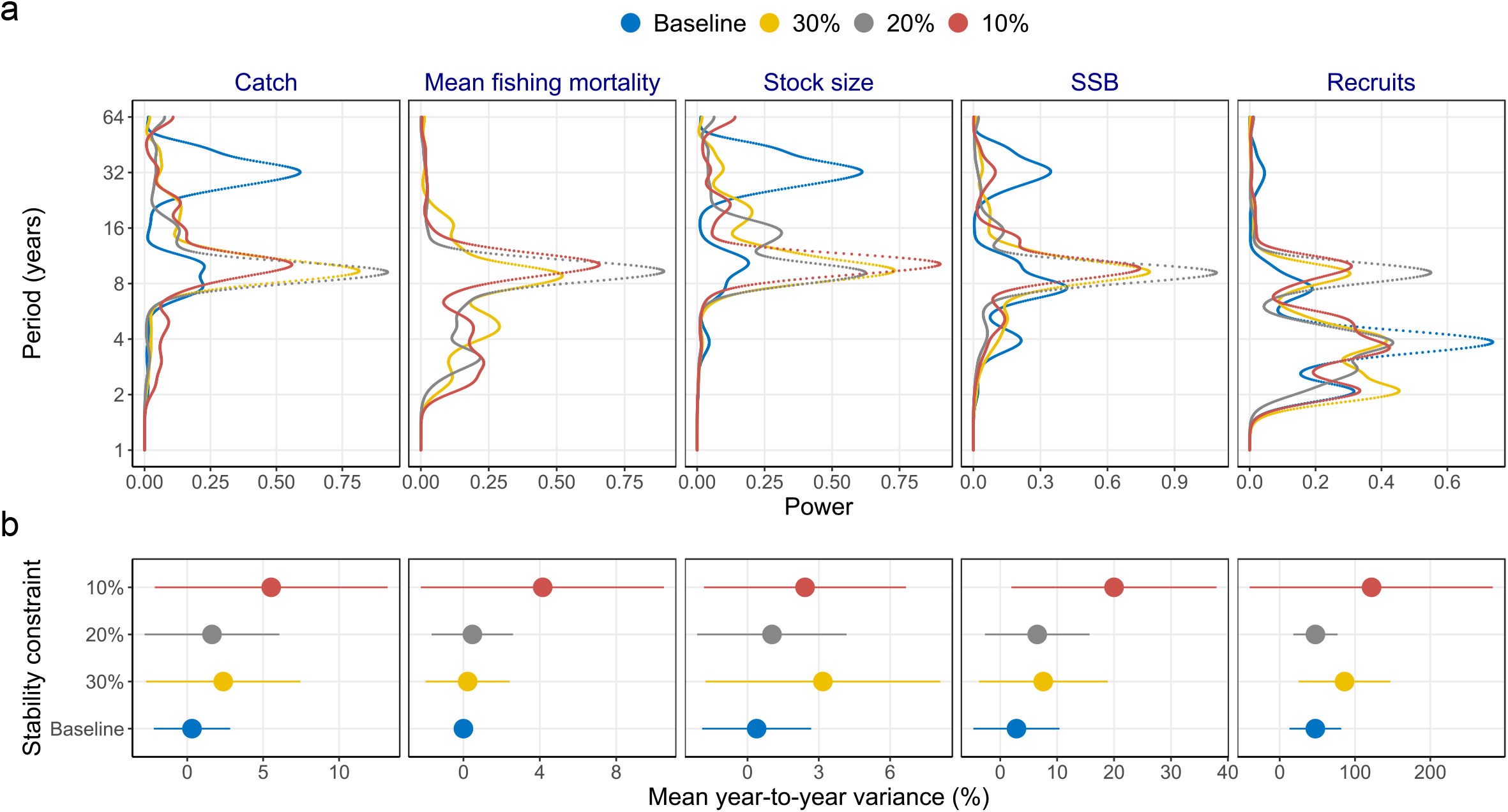
Fluctuations in Northeast Arctic (NEA) cod population behavior and catch projected with a stochastic, multispecies model (STOCOBAR) under scenarios of 0% (baseline), ±30%, ±20%, and ±10% stability constraints. (a) Global (time-averaged) wavelet power spectra of NEA cod catch, mean (5-to 10-year-olds) fishing mortality, harvestable (3-year-olds and older) biomass (stock size), spawner abundance (SSB), and recruits (abundance of 3-year-olds). (b) Mean year-to-year variance (%) in catch, mean fishing mortality, stock size, SSB, and recruits. In a, y-axis is in the logarithms to the base 2. In b, error bars indicate 95% confidence intervals (among-year variation over years 21–100).

### Cod population dynamics and yield stability

Imposed catch stability control (especially 30% or less) increasingly modified the amplitude and periodicity of not only catch forecasts but also cod population behavior (Table 1, Fig. 3, and Appendix 2: Fig. S1 and S2). Under the baseline scenario, wavelet analyses showed modest 32-yr and weak 8-yr cycles in catch, stock size, and SSB (Fig. 3a and Appendix 2: Fig. S2a–c) with low between-year variability (Fig. 3b and Appendix 2: Fig. S1), whereas recruitment showed weaker, high-frequency (2-to 4-yr) cycles (Fig. 3a and Appendix 2: Fig. S2). However, when stability control was introduced, the periodicity of population and harvest fluctuations sharply shifted to higher (~8-yr) frequency cycles with greater amplitudes (by, on average, 57% in catch forecast, 47% in stock size, and 160% in SSB, Fig. 3a and Appendix 2: Fig. S2a–c) and between-year variation (by, on average, 915% in catch forecast, 423% in stock size, 297% in SSB, and 80% in recruitment, respectively, Fig. 3b). The stability control was also imposed increasingly more often as the constraint became stricter (33%, 48%, and 72% of the projection years under 30%, 20%, and 10% constraints, respectively). With variable capelin productivity, mean between-year variation in catch forecast and fishing mortality also rose (ranging from –39% to 166% and –24% to 96%, respectively); catch forecasts fluctuated from 142,000 to 1234,000 t/yr, whereas mean fishing mortality fluctuated from 0.18 to 1.1/yr. These patterns were contrasted with the latest (2016) evaluation of the harvest strategies that was based on assumptions of constant capelin productivity and no temperature dependence, showing that mean between-year variation in catch declines from 26.9% to 12.4% when a 10% constraint was introduced (ICES 2016).

### Population–fishery asynchrony

Temporal fluctuations of cod stock size and mean fishing mortality became increasingly out of phase as the strength of stability control increased (Fig. 4a). Cross-wavelet analyses showed these unsynchronized signals (with ~4–5 yr delays) converging to shorter (~8-yr) cycles with greater amplitudes under constraints of 30% or less (Fig. 4b and Appendix 2: Fig. S3). With being completely out of phase with stock size under a 10% constraint, oscillations in mean fishing mortality became progressively more amplified, peaking in year ~75 before target exploitation rate was substantially reduced by the management model (Fig. 4a).

**Figure 4.**
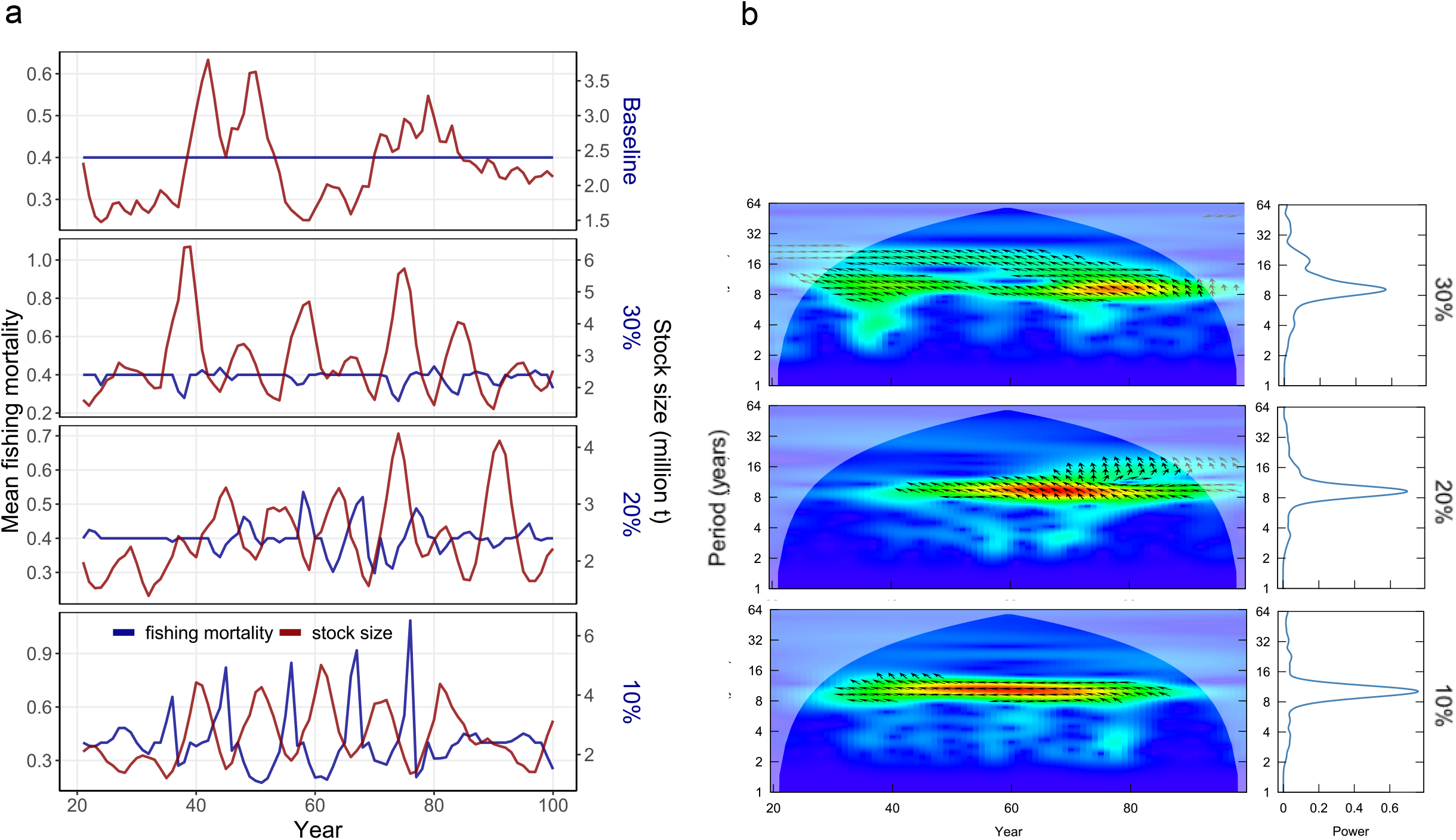
Synchrony of Northeast Arctic (NEA) cod population abundance and fishing pressure projected with a stochastic, multispecies model (STOCOBAR) under scenarios of varying strength of stability constraints. (a) Time series (years 21–100) of NEA cod harvestable (3-year-olds and older) biomass (stock size) and mean (5-to 10-year-olds) fishing mortality under 0% (baseline), ±30%, ±20%, and ±10% constraints. (b) Cross-wavelet power spectra [local (times series, left) and global (time-averaged, right)] between NEA cod stock size and mean fishing mortality under ±30%, ±20%, and ±10% constraints. In b, red areas indicate higher power (intensity of periodicities), blue areas indicate lower power, and white areas indicate regions influenced by edge effects (outside the “cone of influence”) and inferences cannot be made (Cazelles et al. 2014). In b, y-axis is in the logarithms to the base 2; arrows pointing right, left, down, and up indicate the two series being in-phase, the two series being anti-phase, stock size being leading, and mean fishing mortality being leading (respectively).

### Stability-yield tradeoffs

Although average catch forecasts increased (by 10–22%) as the strength of stability control increased (Table 1), overharvest risk became increasingly greater owning primarily to high between-year variability (Fig. 5). With constraints of 30% or more, SSB fell below the lower biological limit threshold, *B_lim_*, with less than 2% of simulations (Fig. 5). By contrast, with constraints of 30% or less, SSB fell below *B_lim_* more frequently (up to 6.5%) because of greater amplitudes resulting from unsynchronized cod population–harvest dynamics (Fig. 4). These risk levels under variable capelin productivity and sea temperature were up to 600x larger than when previously estimated under constant capelin productivity and no temperature-dependence (ICES 2016).

**Figure 5.**
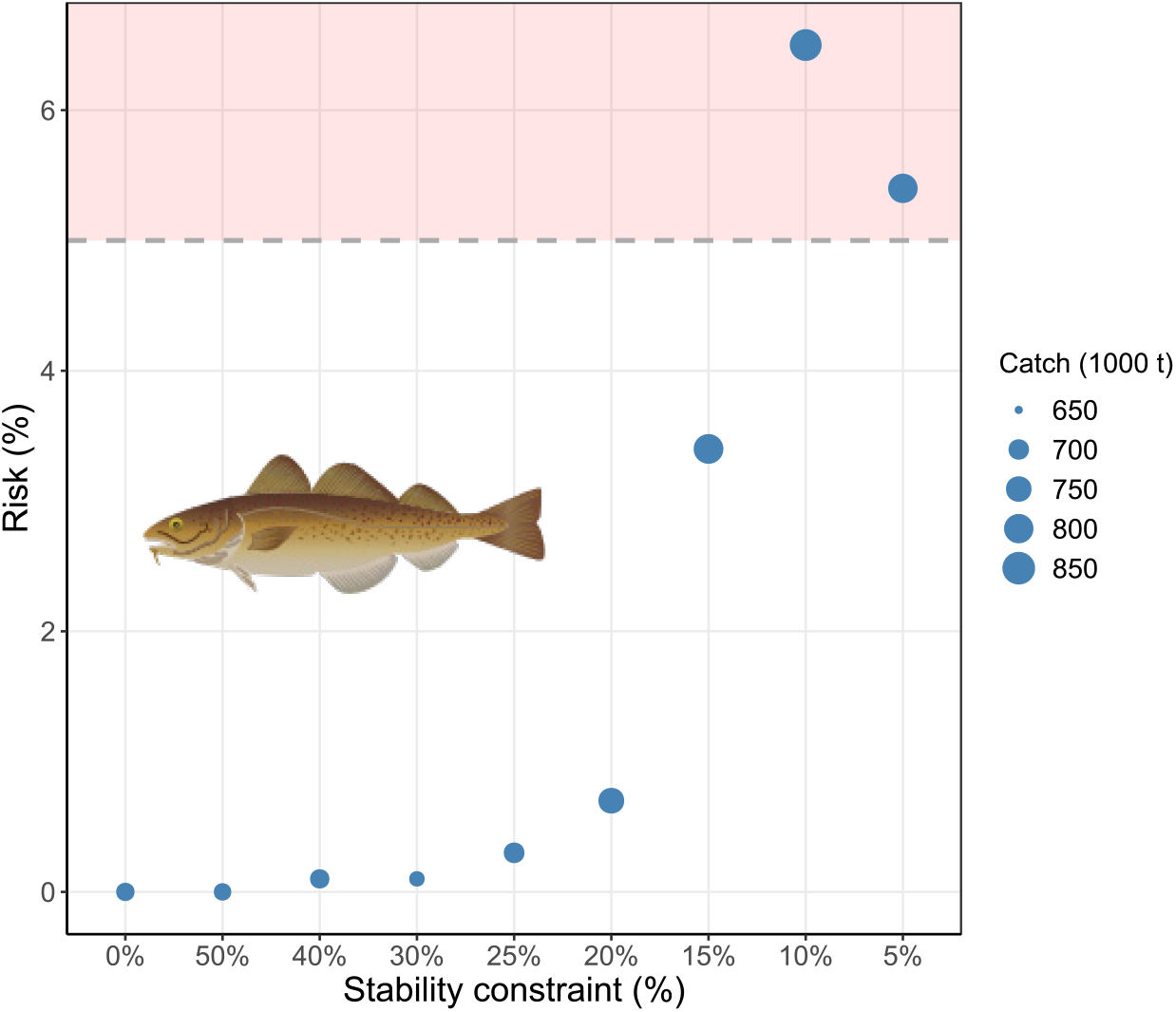
Relationship between stability constraints (±CV%) and risk for Northeast Arctic cod computed from stochastic, multispecies model (STOCOBAR) simulations. Risk is based on the probability of spawner abundance (SSB) falling below the lower biological limit threshold (*B_lim_* = 220,000 t, ICES 2016). A gray horizontal dashed line indicates the risk criterion for reproduction of the population becoming impaired (5%).

## DISCUSSION

We demonstrate that the harvest strategy to suppress short-term yield fluctuation can inadvertently intensify fishing pressure and destabilize population–fishery dynamics of exploited predators if not accounting for uncertainty in how fishing pattern modifies population cycles under fluctuating climate and ocean conditions. Our model-assisted analyses reveal that the cyclicity of stochastic environmental forces (prey abundance and sea temperature) can interactively shape the periodicity and amplitude of predator population and demographic (reproduction and mortality) oscillations, which, in turn, act as a dynamic biological filter (Bjørnstad et al. 2004, Rouyer et al. 2012) to modulate catch fluctuation. A case study with the Northeast Arctic (NEA) cod shows that inter-cohort density-dependent processes such as cannibalism (an additional source of juvenile survival variability, Bjørnstad et al. 1999) can further intensify environmentally driven signals in the population cycle through early life-history events under strict yield stability control. The undesirable (and potentially counterintuitive) consequence of this fishing pattern–enhanced amplitude and shortened periodicity–arises because suppressing between-year variation in adult survival forces fishing pressure to become progressively unsynchronized with population size owing partly to pronounced boom-and-bust cycles of exploited species (Anderson et al. 2008, Frank et al. 2016). Although limiting catch fluctuation (30% or less for NEA cod) theoretically allows, on average, larger yields because of more frequent, larger peaks in stock abundance under fluctuating ocean conditions, the unsynchronized population–fishery dynamics would also increase the probability of overharvest. Incorrect or inappropriate ecological assumptions of predator population cycles when forecasting fishery catches can, thus, overestimate population (and thus yield) stability, thereby underestimating (up to a few orders of magnitude) depletion risk and likely delaying management actions.

### Ecosystem control of exploited population cycles

Myriad ecosystem processes regulating survival of exploited predators interactively influence how given harvest strategies perform in practice (Okamoto et al. 2016). Harvest strategies to regulate fishing pattern unsynchronized with ecosystem carrying capacity, for example, can amplify its variability instead (Link 2017). Our work suggests that suppressing short-term yield fluctuation can accelerate the population cycle of fished species and, in turn, inadvertently elevate variability not only in catch but also in population behavior in stochastically fluctuating environments. In relatively species-poor, high-latitude systems such as the Barents Sea, changes in ecosystem productivity likely propagate further through the food web (less buffering) and may amplify or attenuate year-to-year fluctuation in life history events and population size of fished predators (Bellingeri and Vincenzi 2013). Year-class fluctuation of many fished populations is governed by temporal fluctuation in large-scale climatic and oceanic processes (Cushing 1990) that are mediated through food-web interactions (Lilly et al. 2008). By explicitly accounting for one of the key ecological processes that regulate cod productivity–a density-dependent interplay of cannibalism and food supply, our analyses reveal that the NEA cod population fluctuates with internally and externally regulated, low-frequency (multidecadal) cycles, roughly reflecting capelin productivity cycles when fishing pattern is not forced to suppress yield fluctuation.

Over the past half century, the Barents Sea has shown large fluctuations in productivity (Dalpadado et al. 2012), during which capelin has collapsed three times (Gjøsæter et al. 2009). The capelin collapses subsequently intensified cod cannibalism, weakening cod recruitment strength (Yaragina et al. 2009, Yaragina et al. 2018) and ultimately depleting the cod stock in the late 1980s. Because Barents Sea capelin also supports (in addition to its fishery) other apex predators as a food source, its declines may have triggered a series of trophically mediated processes that exacerbated the cod population depletion, including increased consumption of the young by marine mammals such as harp seals (Bogstad et al. 2015). Declined adult cod survival by overharvest can also release mesopredators from predation (Hjermann et al. 2007), promoting resource competition with and predation on young cod–weakening the ‘cultivation’ effect (Walters and Kitchell 2001). These complex, ecological dynamics that regulate year-class strength of fished populations can have both stabilizing and destabilizing effects under variable fishing pressure and ecosystem productivity (Anderson et al. 2008) and need to be accounted for in designing and evaluating management measures.

### Managing harvest stability of predators with transient cycles

Imposing yield stability may inadvertently amplify variability of exploited species by modifying natural population cycles owing to various sources of uncertainty often overlooked in assessments (Botsford et al. 1997). Nearly three-quarters of assessed fish stocks worldwide (200+) have shown irregular temporal shifts in productivity regime (Vert-pre et al. 2013); imposing stability control without accounting for ecological uncertainty such as fluctuation in forage species productivity may increase overharvest and depletion risks of fished predators. In the case of NEA cod, analyses indicate that an attempt to dampen the amplitude of yield fluctuation instead amplifies its sensitivity to environmental forces with stronger signals–sea temperature in this study–and shorten population cycle, thereby forcing the exploited predator to become increasingly unsynchronized with (and, in effect, inadvertently intensifying) fishing pressure. Further, the periodicity of population and catch fluctuations progressively shifts toward higher frequencies that reflect life-history traits such as the dominant age groups of spawners (~6-to 8-year-olds for NEA cod, Appendix 1: Fig. 1 and Rouyer et al. 2011) as fishing pressure becomes intensified (through stricter stability control), a pattern predicted by theory (Bjørnstad et al. 2004, Worden et al. 2010). Because industrial capture fisheries often selectively remove larger and older individuals, biased reduction in adult survival by fishing can further modify demography and population cycle by truncating age structure (juvenescence, Lindegren et al. 2013, Okamoto et al. 2016). As a result, intensely fished populations become progressively more recruitment-dependent, eroding their resilience to environmental and demographic stochasticity (Minto et al. 2008, Hidalgo et al. 2011), confounding efforts to sustain human exploitation while conserving biological diversity in the ecosystems.

Although large short-term variation in catch is often disfavored by harvesters and seafood processors, oscillatory patterns in population cycle of exploited predators may provide valuable information about the population status such as its resilience to changes in external forcing (Anderies 2015). Magnified variance may signal impending internal structural shifts in nature including fished populations (Brock and Carpenter 2010). The analysis of the extensive database on global commercial marine fisheries compiled by the United Nations Food and Agriculture Organization (FAO) suggests that imposed catch stability can hide early warning signals that may help prevent population collapses–a potential long-term consequence of suppressing short-term yield fluctuation (Mullon et al. 2005, Carpenter et al. 2015). Stability in exploitation of living marine resources may, thus, need to come from promoting population resilience (restoration of age and size structure, for example) rather than imposing apparent stability (Anderson et al. 2008, Carpenter et al. 2015).

Our case study with the exploited Arctic piscivore–planktivore complex illustrates that the harvest strategy to suppress yield fluctuation–a widely implemented policy tool to achieve socioeconomic goals–can bring about counterintuitive, adverse outcomes if not accounting for fluctuating ocean conditions. This harvest policy can inadvertently destabilize population–fishery dynamics of exploited species as it forces fishing pattern to become unsynchronized with the cyclicity of population size and demography under environmental stochasticity. Because our model did not account for other sources of uncertainty such as observation errors and model structural errors (for example, no predation feedback from adult cod to capelin), year-to-year population and harvest fluctuations in simulations are likely conservative estimates. Nevertheless, the findings highlight broader implications of ecological uncertainty when designing and analyzing harvest policies for ecologically connected populations. Evaluating a policy without accounting for variability in key ecosystem processes–common practice in assessments of exploited species–may fundamentally reshape population structure and behavior, thereby underestimating the risk of population decline and possibly extinction (Gårdmark et al. 2013). With expected changes (both mean and variance) in climatic fluctuation and, in turn, ocean productivity (Wassmann et al. 2011, Cohen et al. 2014, Lewis et al. 2020), adapting ecosystem approaches tailored to specific contexts (such as use of models of intermediate complexity for ecosystem, Plagányi et al. 2014), as demonstrated here, will safeguard against such risk and help sustainably manage marine living resources while securing socioeconomic stability.

## ACKNOWLEDGEMENTS

We thank anonymous reviewers for comments on earlier versions of the manuscript. Some figures use images from the IAN Symbols, courtesy of the Integration and Application Network, University of Maryland Center for Environmental Science (ian.umces.edu/symbols/). This project was partially funded by Institute of Marine Research’s REDUS (Reduced Uncertainty in Stock Assessments) project.

