## Appendix 1 for "Trade-offs of managing Arctic predator harvest instability in fluctuating oceans"

**Appendix S1.** Additional results of Northeast Arctic cod population dynamics projected with a stochastic, multispecies model (STOCOBAR) under the baseline scenario.

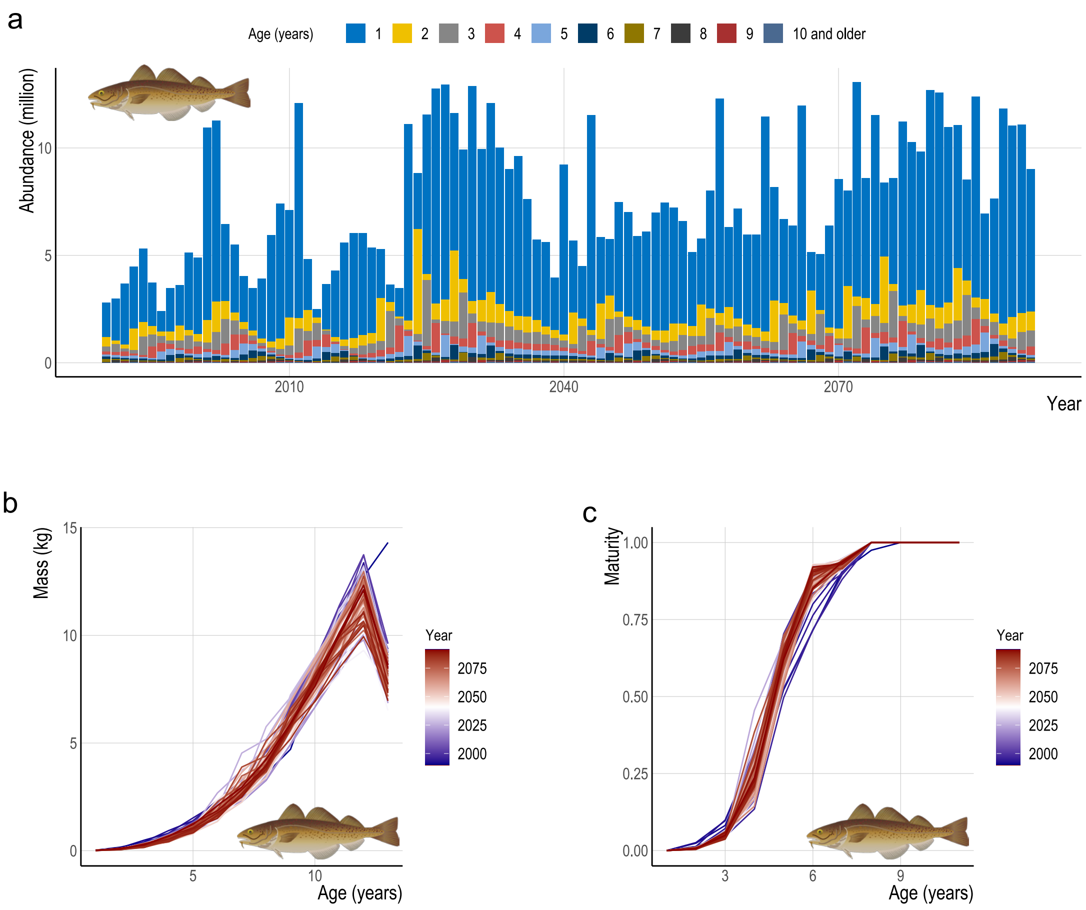


Figure S1. Northeast Arctic cod population demographic state variables projected with a stochastic, multispecies model (STOCOBAR) under the baseline scenarios. (a) Time series of age-specific abundance (millions). (b) Among-year variability in relationships between age and mass (kg). (c) Among-year variability in relationships between age and maturity rate (proportion of adult abundance).

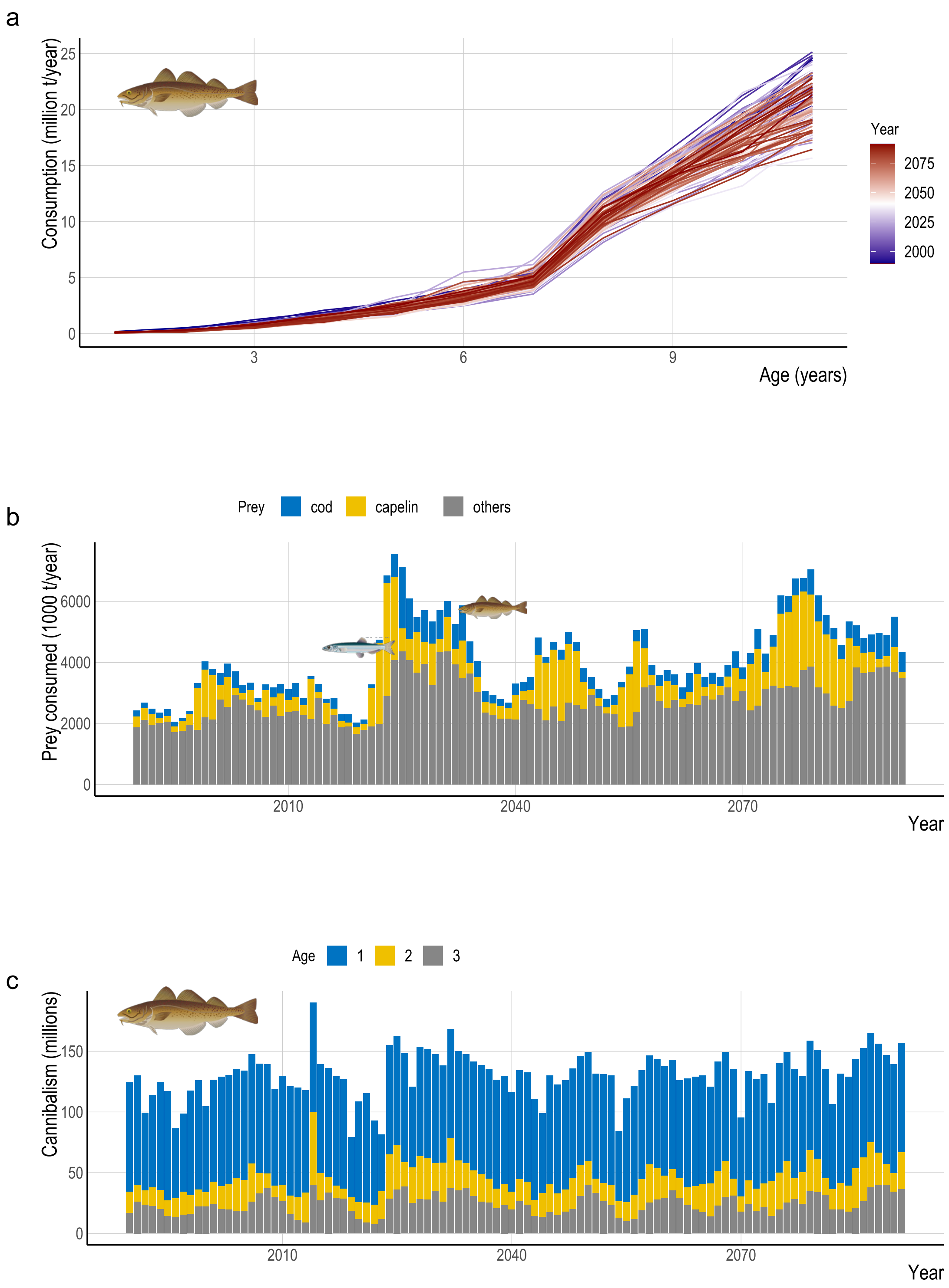


Figure S2. Northeast Arctic cod population trophic dynamics projected with a stochastic, multispecies model (STOCOBAR) under the baseline scenarios. (a) Among-year variability in relationships between age and annual food consumption (million t). (b) Time series of total annual prey consumption (1000 t). (c) Time series of age-specific cannibalism (millions).

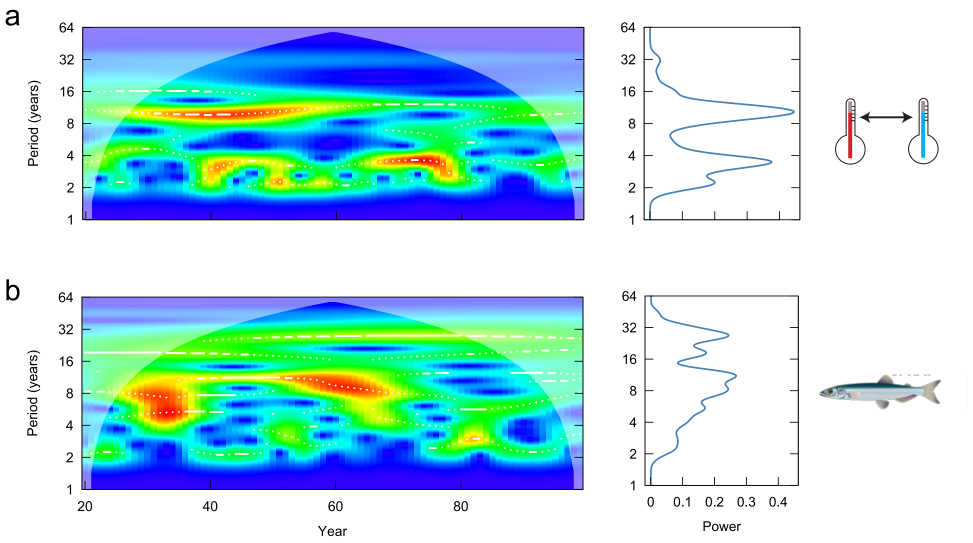


Figure S3. Wavelet power spectra [local (times series, left) and global (time-averaged, right)] computed for (a) modeled sea temperature in the Kola section of the Barents Sea and (b) modeled capelin biomass used in stochastic, multispecies model (STOCOBAR) simulations. In local wavelet spectra, color gradient indicates red areas being higher power (intensity of periodicities) to blue areas being lower power; white areas indicate regions influenced by edge effects (outside the "cone of influence") and inferences cannot be made. Y-axis is in the logarithms to the base 2. Artwork: Courtesy of the Integration and Application Network, University of Maryland Center for Environmental Science (ian.umces.edu/symbols/).
