## Appendix 2 for "Trade-offs of managing Arctic predator harvest instability in fluctuating oceans"

**Appendix S2.** Additional results of Northeast Arctic cod population dynamics projected with a stochastic, multispecies model (STOCOBAR) under scenarios of stability constraints.

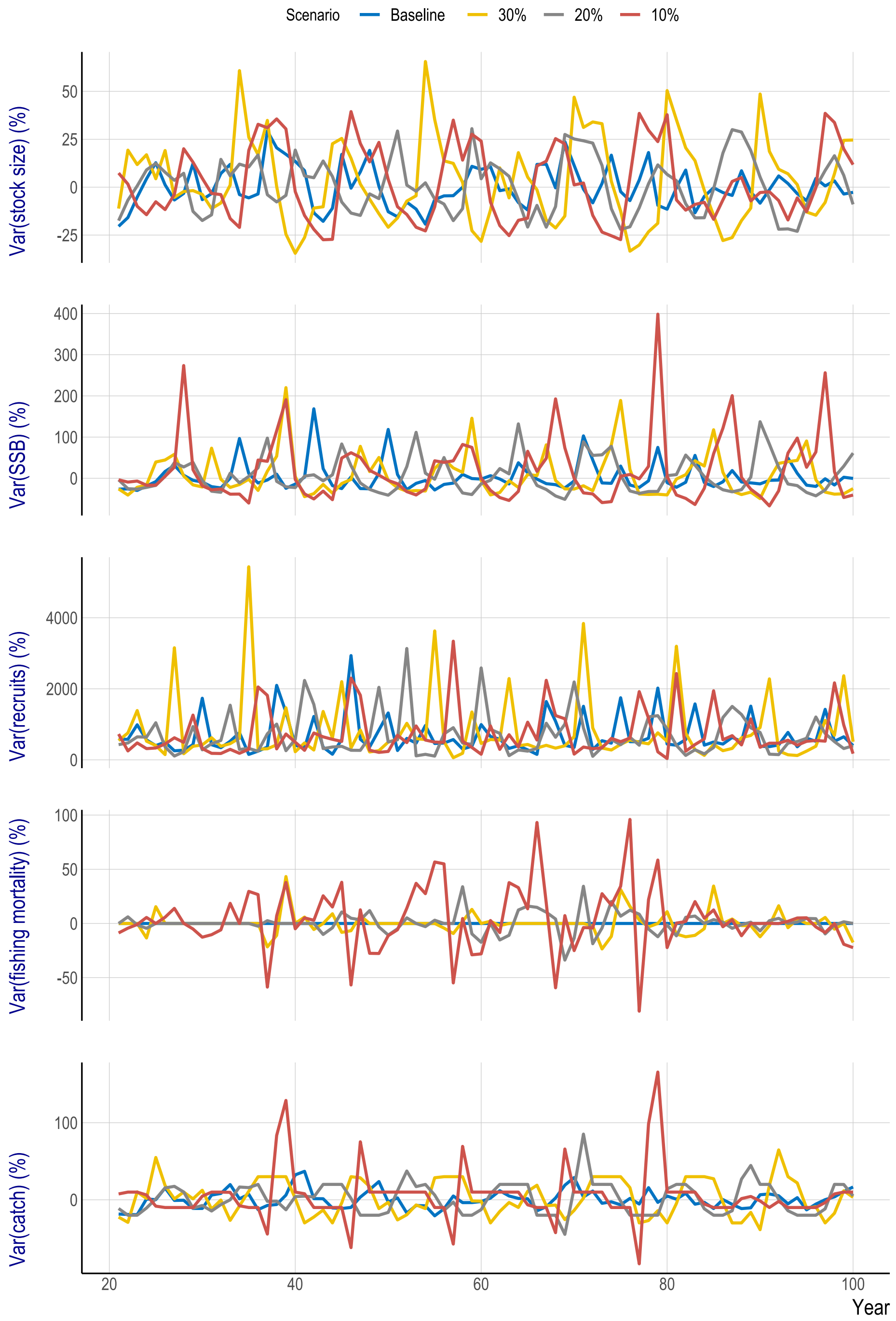


Figure S1. Time series (years 21–100) of between-year variance (%) in total harvestable (3-year-olds and older) biomass (stock size), spawner abundance (SSB), recruits (abundance of 3-year-olds), and mean (5- to 10- year-olds) fishing mortality, and annual catch forecasts of Northeast Arctic cod projected with a stochastic, multispecies model (STOCOBAR) under scenarios of 0% (baseline), ±30%, ±20%, and ±10% constraints.

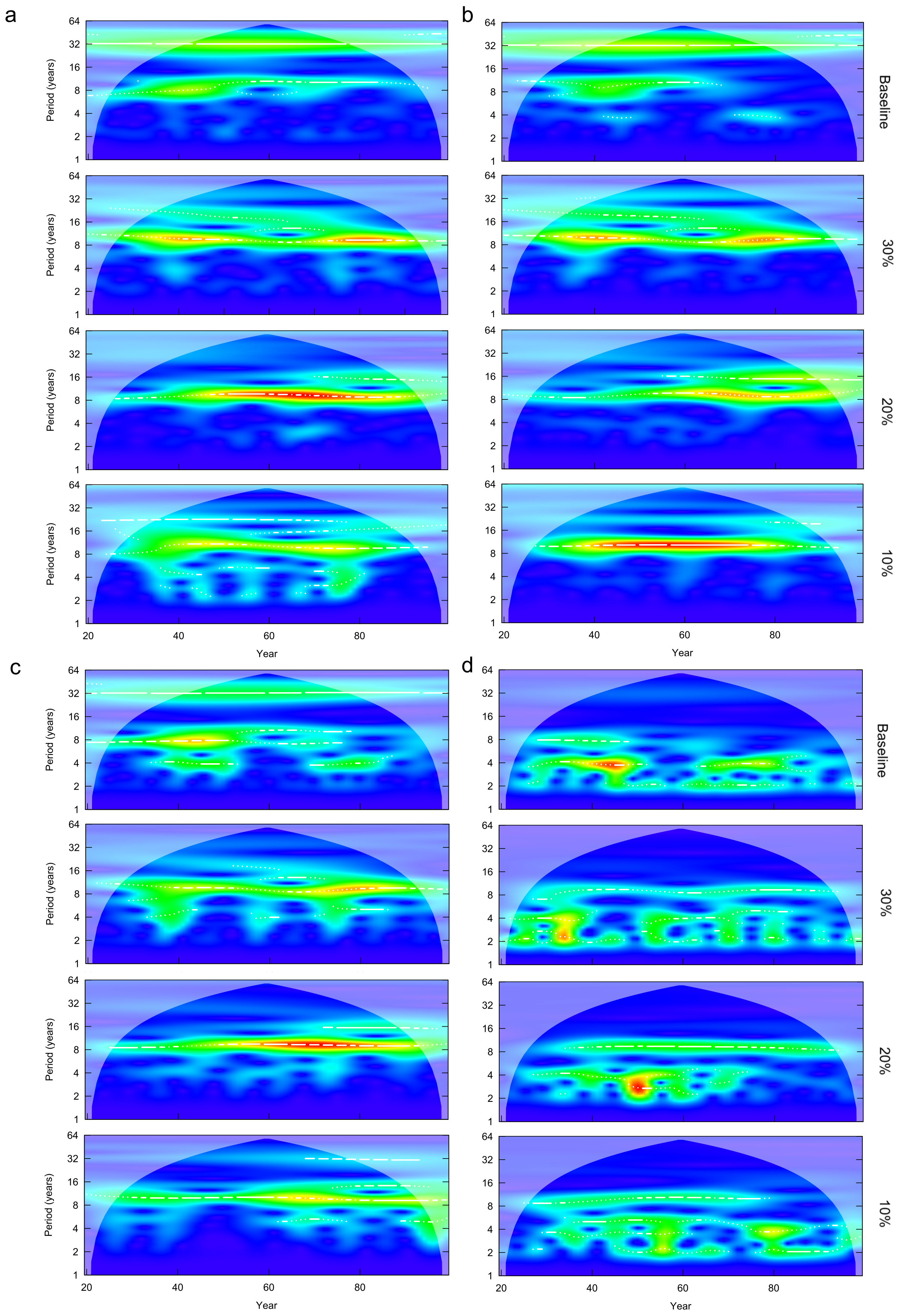
a
b
c
d
Baseline
30%
20%
10%
Period (years)
64
32
16
8
4
2
1
Year
20
40
60
80

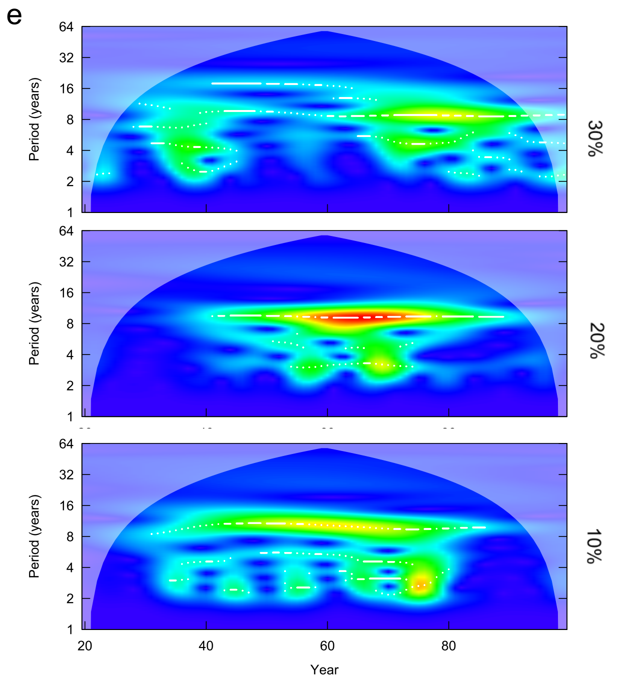


Figure S2. Wavelet power spectra computed for (a) annual catch forecasts, (b) total harvestable (3-year-olds and older) biomass (stock size), (c) spawner abundance (SSB), (d) recruits (abundance of 3-year-olds), and (e) mean (5- to 10- year-olds) fishing mortality of Northeast Arctic cod projected with a stochastic, multispecies model (STOCOBAR) under scenarios of 0% (baseline), ±30%, ±20%, and ±10% constraints. Color gradients indicate red areas being higher power (intensity of periodicities) to blue areas being lower power; white areas indicate regions influenced by edge effects (outside the "cone of influence") and inferences cannot be made. Y-axis is in the logarithms to the base 2.

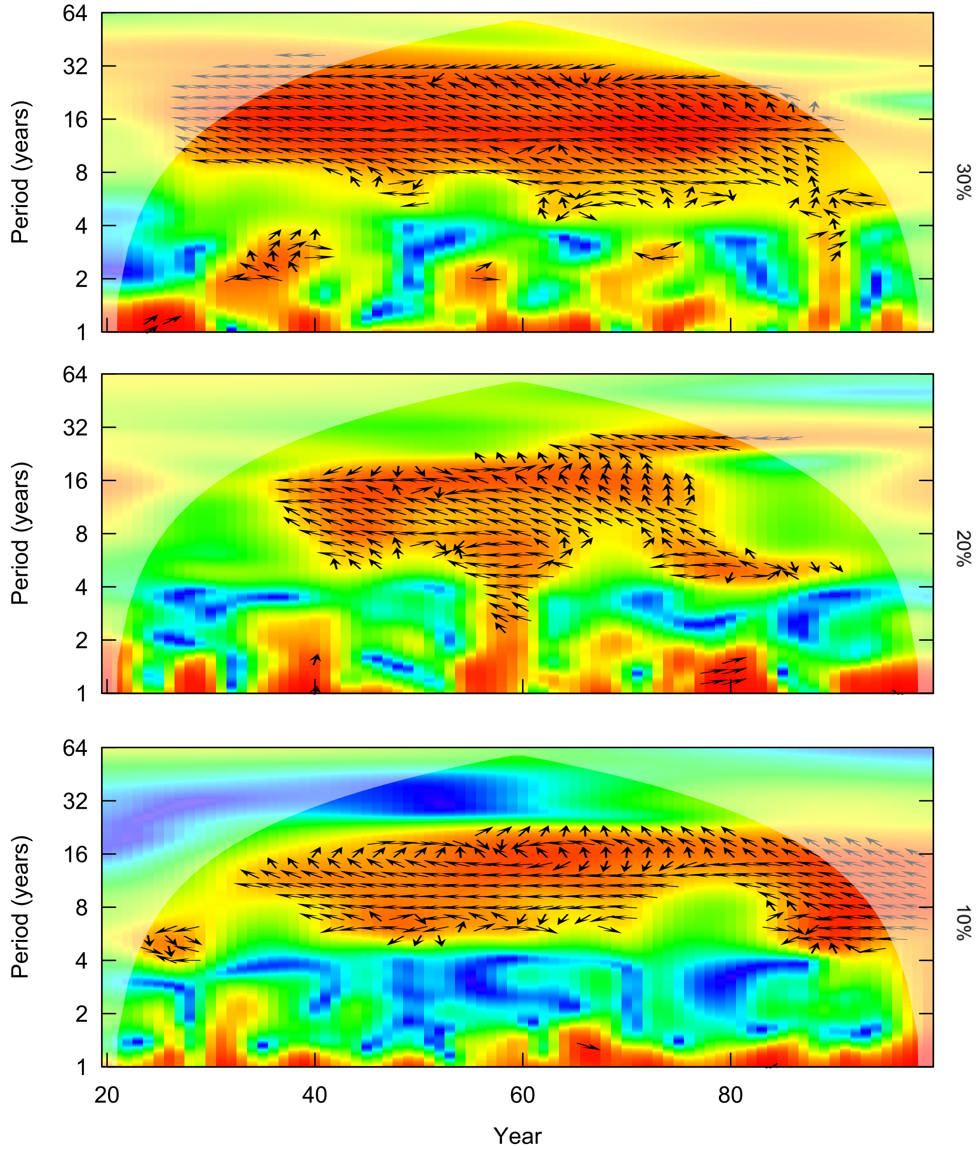


Figure S3. Wavelet coherence spectra computed for total harvestable (3-year-olds and older) biomass (stock size) and mean (5- to 10- year-olds) fishing mortality of Northeast Arctic cod projected with a stochastic, multispecies model (STOCOBAR) under scenarios of ±30%, ±20%, and ±10% constraints. Color gradients indicate red areas being higher power (intensity of periodicities) to blue areas being lower power; white areas indicate regions influenced by edge effects (outside the "cone of influence") and inferences cannot be made. Y-axis is in the logarithms to the base 2; arrows pointing right, left, down, and up indicate the two series being in-phase, the two series being anti-phase, stock size being leading, and mean fishing mortality being leading (respectively).
